# Insulin resistance in cavefish as an adaptation to a nutrient-limited environment

**DOI:** 10.1101/179069

**Authors:** Ariel Aspiras, Misty Riddle, Karin Gaudenz, Robert Peuß, Jenny Sung, Brian Martineau, Megan Peavey, Andrew Box, Julius A. Tabin, Suzanne McGaugh, Richard Borowsky, Clifford J. Tabin, Nicolas Rohner

## Abstract

Periodic food shortage is one of the biggest challenges organisms face in natural habitats. How animals cope with nutrient limited conditions is an active area of study, of particular relevance in the context of the current increasing destabilization of global climate. Caves represent an extreme setting where animals have adapted to nutrient-limited conditions, as most cave environments lack a primary energy source. Here we show that cave-adapted populations of the Mexican Tetra, *Astyanax mexicanus,* have dysregulated blood glucose homeostasis and are insulin resistant compared to the river-adapted population. We found that multiple cave populations carry a mutation in the insulin receptor that leads to decreased insulin binding *in vitro*. Surface/cave hybrid fish carrying the allele weigh more than non-carriers, and zebrafish genetically engineered to carry the mutation similarly have increased body weight and insulin resistance. Higher bodyweight may be advantageous in the cave as a strategy to cope with infrequent food. In humans, the identical mutation in the insulin receptor leads to a severe form of insulin resistance and dramatically reduced life-span. However, cavefish have a similar lifespan to surface fish (of greater than fourteen years) and do not accumulate advanced glycated end products (AGEs) in the blood that are typically associated with progression of diabetes-associated pathologies. Our findings raise the intriguing hypothesis that cavefish have acquired compensatory mechanisms that allow them to circumvent the typical negative effects associated with failure to regulate blood glucose.

---

Animals have evolved a range of physiological strategies to survive periods when nutrients are limited or absent, such as hibernation, diapause, torpor, and estivation^1^. Such strategies serve as a bridge to sustain the animal through a period of deprivation to a time of greater nutrient abundance. Cave-dwelling animals, in particular, must withstand long periods of nutrient limitation, as caves lack photosynthesis-driven primary producers, and hence cave food chains are dependent on energy input from external sources, such as seasonal floods or bat droppings^2^. To thrive under such harsh and unpredictable conditions, cave animals must evolve unique metabolic strategies that are currently not well understood.

The Mexican Tetra, *Astyanax mexicanus*, is a particularly useful model to study the genetic basis of metabolic adaptation. This species consists of a river-dwelling (surface) population and multiple cave-dwelling (cave) populations (Fig. 1a) that experience dramatically different nutrient availablility. In this study, we focused on three cavefish populations (named for the caves they inhabit: Tinaja, Pachón, and Molino) that have evolved independently from two different stocks of surface fish that invaded caves millions of years ago^3^. Tinaja and Pachón cavefish likely originated from a more ancient surface population than the Molino population^4^. More importantly, the surface morphs and all cave morphs are inter-fertile, allowing for genetic studies both in the wild and under laboratory conditions^5,6,7^.

**Figure 1.**
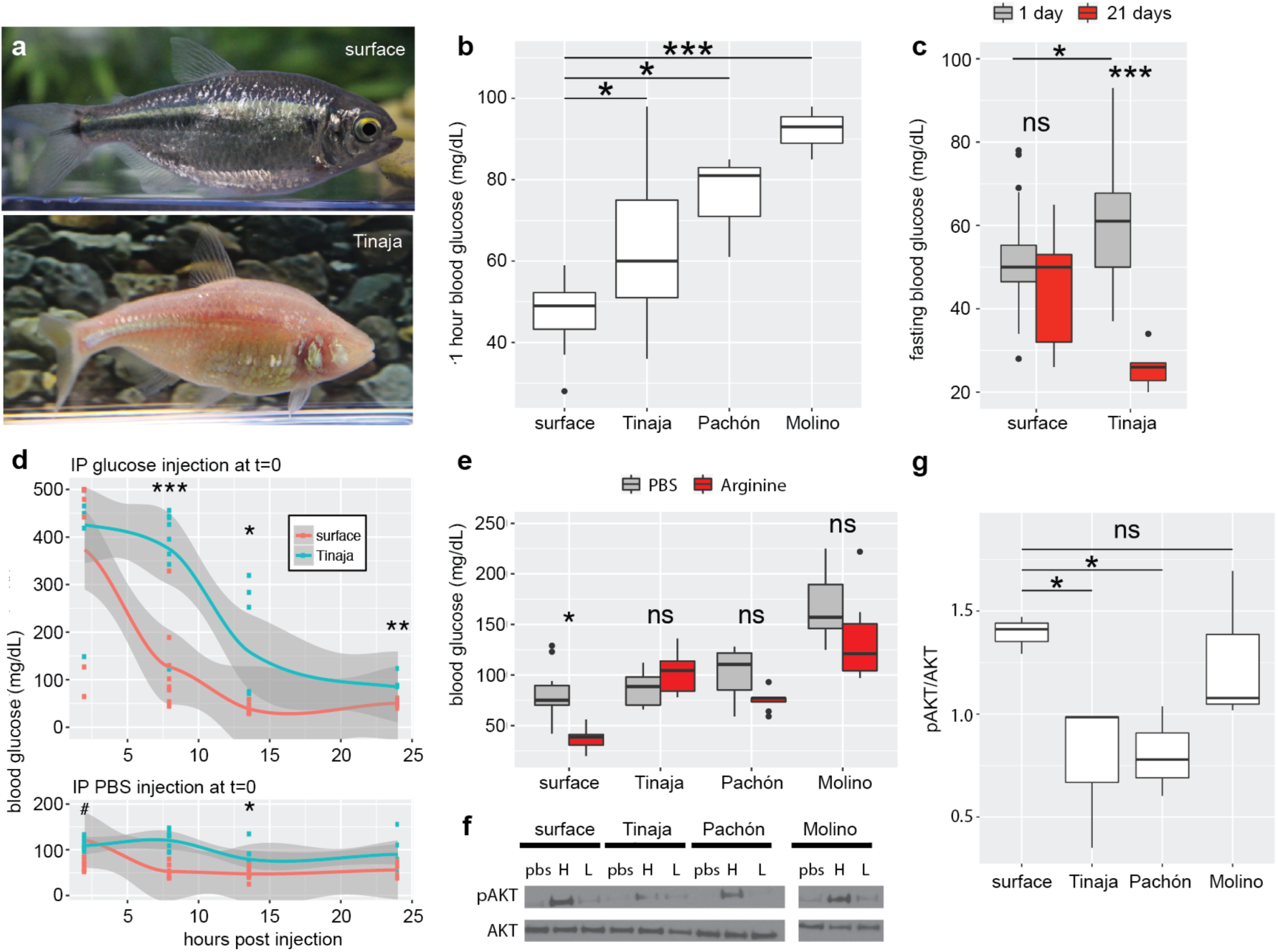
Altered glucose homeostasis in cave-adapted *Astyanax mexicanus* populations (Tinaja, Pachón, Molino) **a**, Photograph of surface fish and Tinaja cavefish of the same species, *Astyanax mexicanus*. **b**, Blood glucose levels one hour after eating in surface fish compared to three independently-evolved cavefish populations (n=10, 13, 3, 3 respectively). **c**, Blood glucose level in surface and cave (Tinaja) populations after 1 day (n=28 fish per population) versus 21 days (n=9 fish per population) without food. **d**, Glucose tolerance test. Blood glucose level after IP injection of glucose (top) or PBS (bottom). Data points represent values for individual fish and grey shade indicates 95% confidence interval for polynomial regression. (# outlier for Tinaja with blood glucose level of 500 mg/dL not shown on graph). **e**, Blood glucose level 5 hours after inducing both glucagon and insulin by IP injection of arginine (n=10 fish per population and condition). **f,** Representative western blot showing cell lysates probed with pAKT (ser473) and AKT antibodies. Lysates were produced from skeletal muscle treated *ex vivo* with pbs, high (1x), or low (0.1x) level of recombinant insulin (x=9.5-11.5 µg/mL insulin). **g**, Quantification of bands by densitometry of the highest concentration treatment (n=3 fish per population). Significance codes from one-way ANOVA with HSD post hoc test, ns p>0.05, *p<0.05, **p<0.005, ***p<0.0005.

**Figure 2:**
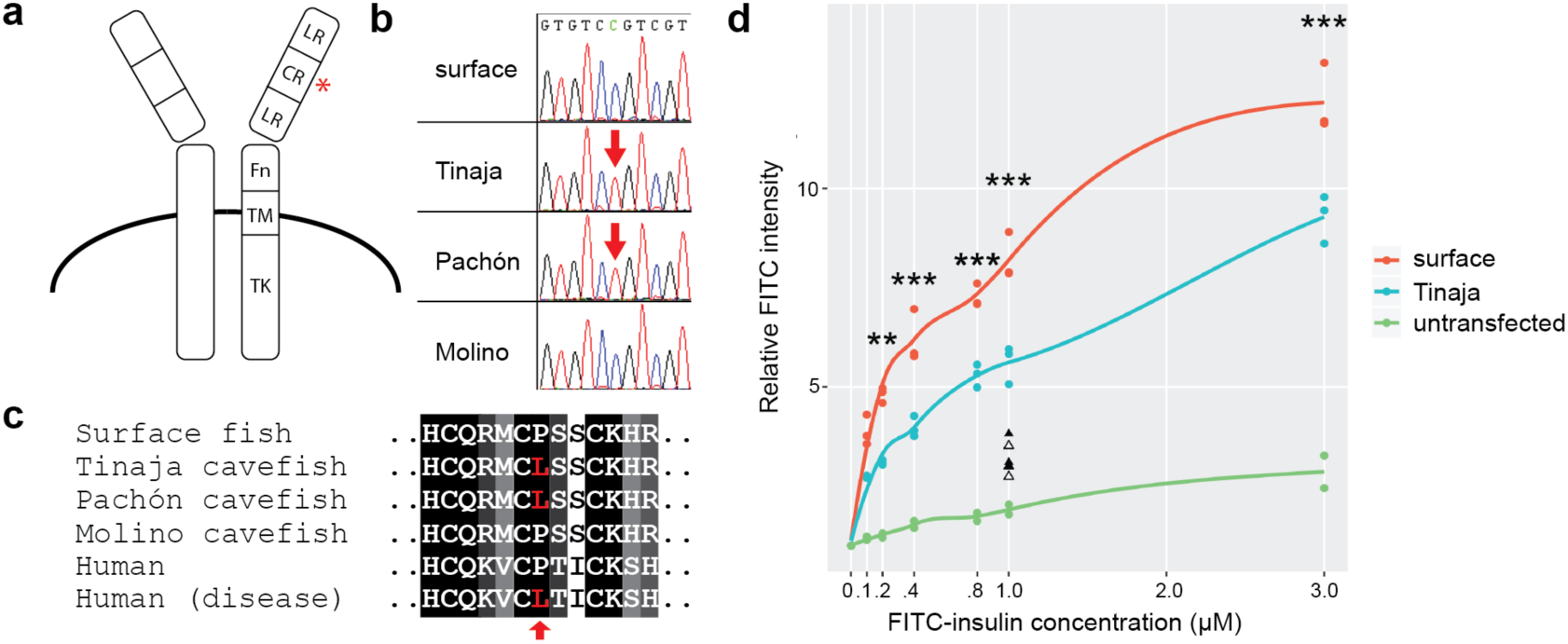
Coding mutation in the cavefish insulin receptor leads to decreased insulin binding. **a**, Schematic representation of the insulin receptor and main domains (adapted from^20^. The red asterisk depicts position of P211L mutation in cavefish. LR, leucine-rich repeats; CR, cysteine-rich domain; Fn, fibronectin type III domain; TM, transmembrane domain; TK, tyrosine-kinase domain. **b**, Sequence chromatogram of the mutation present in Pachón and Tinaja cavefish populations. **c**, Amino acid alignment of the P211L mutation and neighboring amino acids of the insulin receptor in *Astyanax mexicanus* and healthy humans and patients with Rabson-Mendelhall syndrome. **d**, Relative FITC intensity of HEK293T (Flp-In-293) cells stably transfected with FLAG-tagged surface fish or Tinaja cavefish insulin receptors and incubated with indicated concentration of FITC-labeled insulin. Each point represents mean FITC intensity of >2,500 live cells normalized to the mean intensity of untreated cells. Lines represent results from local polynomial regression fitting. Triangles represent data from competitive binding assay: surface (filled) or Tinaja (unfilled) Flp-In-293 cells were incubated with the addition of 10µM unlabeled insulin. Significance codes are from one-way ANOVA (between surface and Tinaja) with Tukey’s HSD post hoc test, *p<0.05, **p<0.005, ***p<0.0005.

We have previously shown that cavefish are starvation resistant compared to surface fish and lose a smaller fraction of their bodyweight after two months without food ^8^. Several factors have been identified that contribute to starvation resistance, including a reduction in metabolic circadian rhythm^9^, a general decrease in the metabolic rate^10^, and elevated body fat levels that serve as an energy dense reserve^8,11^. However, the genetic changes underlying these extreme metabolic adaptations remain largely unknown.

A critical aspect of metabolic homeostasis is the regulation of blood glucose^12^. We compared the blood glucose levels of lab-raised surface fish with lab-raised cavefish (Tinaja, Pachón, Molino) and found that one hour after eating, all three cave populations had significantly higher blood glucose (63, 76, 92 vs 51mg/dL respectively, Fig. 1b). We further investigated the dynamics of glucose homeostasis during fasting by comparing surface and Tinaja populations. While Tinaja cavefish had significantly higher blood glucose levels after 24 hours of fasting (60 vs 52mg/dL, p=0.05, Fig. 1b), they were unable to maintain glucose homeostasis long-term, reaching as low as 22mg/dL after 21 days without food (mean 26mg/dL, Fig. 1c). In contrast, surface fish blood glucose did not change significantly over the same fasting period (52 vs. 45 mg/dL, p=0.49). Our findings suggest that surface fish have tighter control of blood glucose levels.

To further test this hypothesis, we compared the acute control of glucose homeostasis using a glucose tolerance test (Fig. 1d). We injected glucose into the intraperitoneal (IP) cavity of both surface fish and Tinaja cavefish, transiently raising blood glucose levels to over 400 mg/dL in most fish (Fig. 1d). After eight hours, the blood glucose levels of the surface fish returned to the same levels as the PBS-injected controls (mean 126 vs 120mg/dL, Fig. 1d), while blood glucose levels remained highly elevated in Tinaja cavefish, (mean 374mg/dL, p<0.0005, Fig. 1d). Tinaja cavefish blood glucose levels decreased over time, but remained elevated compared to surface at both 14hrs (158 vs. 38mg/dL, p<0.05, Fig. 1d) and 24hrs (85 vs. 51mg/dL, p<0.005, Fig. 1d) after glucose injection. These results suggest that cavefish have impaired glucose clearance.

Glucose homeostasis requires the balanced release of the hormones insulin and glucagon that signal to peripheral tissues to absorb glucose from the blood or produce glucose from stored glycogen^13^. We compared the levels and actions of these hormones between surface and Tinaja fish to gain further insight into their differences in glucose regulation. We did not detect a difference in the number of cells producing glucagon (54 vs 50, p=0.678) or insulin (54 vs 52, p=0.275) in 10-day old fish, (n=5 fish per population, Extended Data Fig. 1) and similarly we did not detect a difference in circulating glucagon levels in adult fish (Extended Data Fig. 2). Circulating insulin levels tended to be higher in Tinaja cavefish, but the results were not significant (n=24 fish per population, 2 age groups, and 3 samples per fish, Extended Data Fig. 3). Nonetheless, we found evidence of diminished insulin response in cavefish: we injected arginine into surface fish and cavefish to stimulate the simultaneous release of glucagon and insulin^14^ and observed that while surface fish had a significant decrease in blood glucose level (80 vs 38mg/dL, p=0.006), Tinaja, Pachón, and Molino blood glucose did not change (Fig. 1e). In addition, we found that injection of recombinant human insulin caused a significant drop in blood glucose after 60 minutes in Surface fish but not in Tinaja cavefish (Extended data Fig.4). Our combinded observations that glucagon and insulin levels do not differ between surface and cavefish (Extended Data Fig. 2), and that cavefish do not decrease blood glucose levels in repsonse to arginine or insulin (Fig, 1e, Extended data Fig. 4) suggest that cavefish may be insulin resistant.

**Figure 3:**
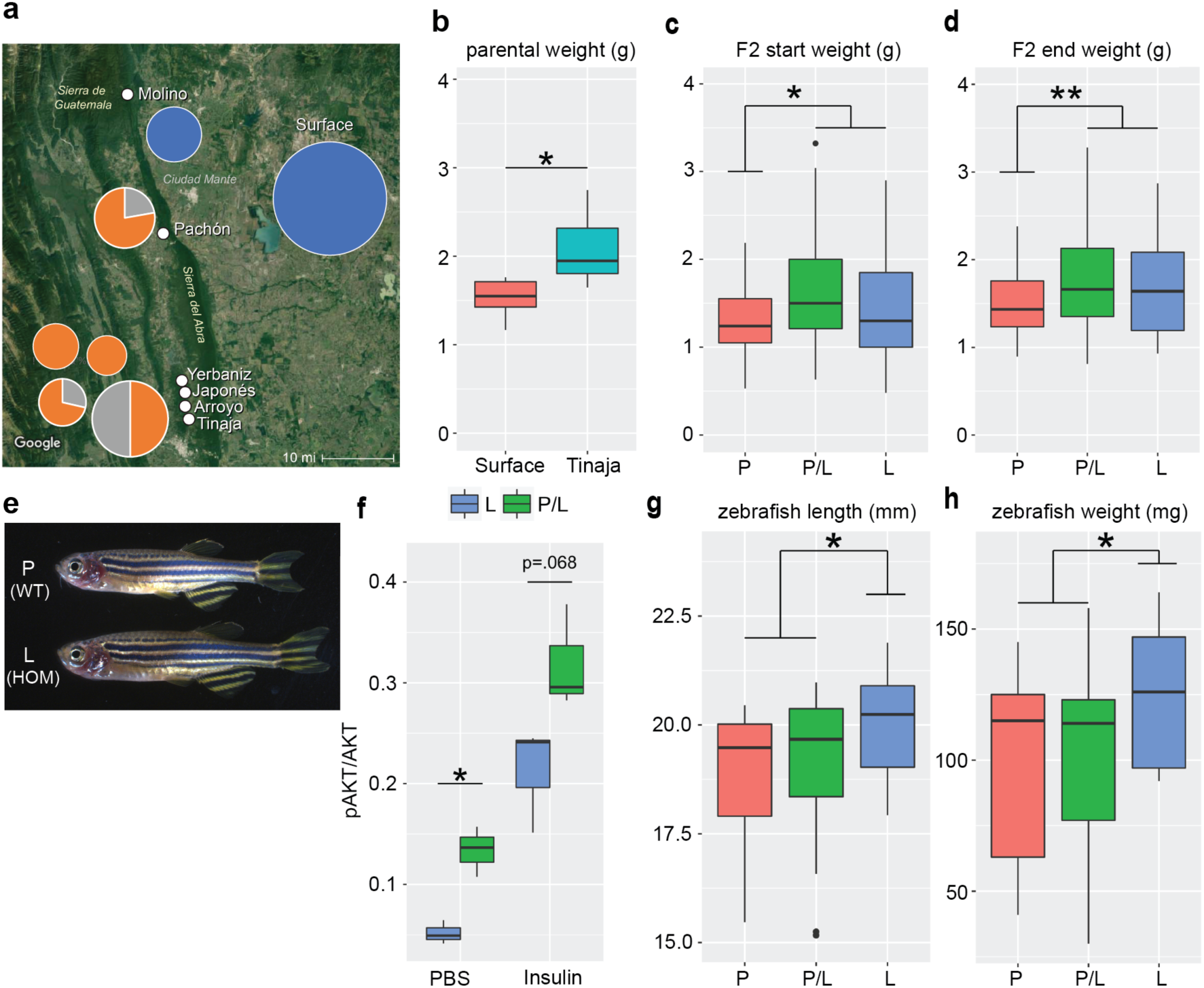
The P211L *insra* allele shows signs of selection in cave environments and is associated with higher body weight in surface/cave hybrids. **a,** Genotyping of the *insra* gene for the presence of the derived allele (P211L) in wild-caught samples. Pie charts indicate percentage of fish with the surface allele (blue), cave allele (orange), or both (grey), and size of pie chart roughly indicates number of fish genotyped (Molino n= 8, Surface n= 71, Pachón n= 9, Yerbaniz n =8, Japonés n = 5, Arroyo n= 7, Tinaja n=14). The cave allele is absent in all the wild-caught surface fish and Molino cavefish (new lineage). In all sampled cavefish populations from the old lineage, the P211L mutation was present. Tinaja, Yerbaniz, Japonés, and Arroyo are geographically close and believed to represent a single invasion event; Pachón represents an independent invasion^26^. The deviation from Hardy-Weinberg equilibrium suggests that selection is acting on the locus favoring the presence of the mutation in the cave environments. Map source: Imagery ©2017 Landsat/Copernicus, Map data ©2017 Google, INEGI **b**, Boxplot showing weight of Tinaja males (n=6) and surface males (n=5) on a nutrient limited diet. **c**, Weight of 18-month-old F2 male Tinaja/surface hybrids sorted for presence of the P211L mutation (P-homozygous surface (n=22), L-homozygous cave (n=27), P/L heterozygotes (n=53)). Mean weight of F2 fish homozygous for the surface allele (1.28 gram; n=22) is significantly less than F2 fish carrying at least one of the cave alleles (1.63 gram; n=80; *p=0.006). **d**, Weight of the same F2 fish after four months on a limited nutrient diet. Mean weight of F2 fish homozygous for the surface allele (1.47 gram; n=22) is significantly less than F2 fish carrying at least one of the cave alleles (1.82 gram; n=80; **p=0.004). Absolute weight change (not shown, surface allele: 0.193 gram, cave allele: 0.185 gram) or percent weight change (surface allele: 20.5%, cave allele 17.3%) is not significant (p=0.88, and p=0.67 respectively). **e,** Representative pictures of a WT sibling and a homozygous P211L zebrafish mutant. **f**, Ratio of pAKT/AKT in adult zebrafish skeletal muscle treated *ex vivo* with insulin or PBS. The ratio of pAKT/AKT is signficcantly less in zebrafish homozygous for the P211L mutation compared to heterozygous fish (n=3 fish per genotype and condition). Significance codes are from two-tailed students t-test, *p<0.05, **p<0.005. **g, h,** Length and weight of 55dpf zebrafish sorted by presence of the P211L mutation (n=11(L), 13(P), 22(P/L)).

**Figure 4:**
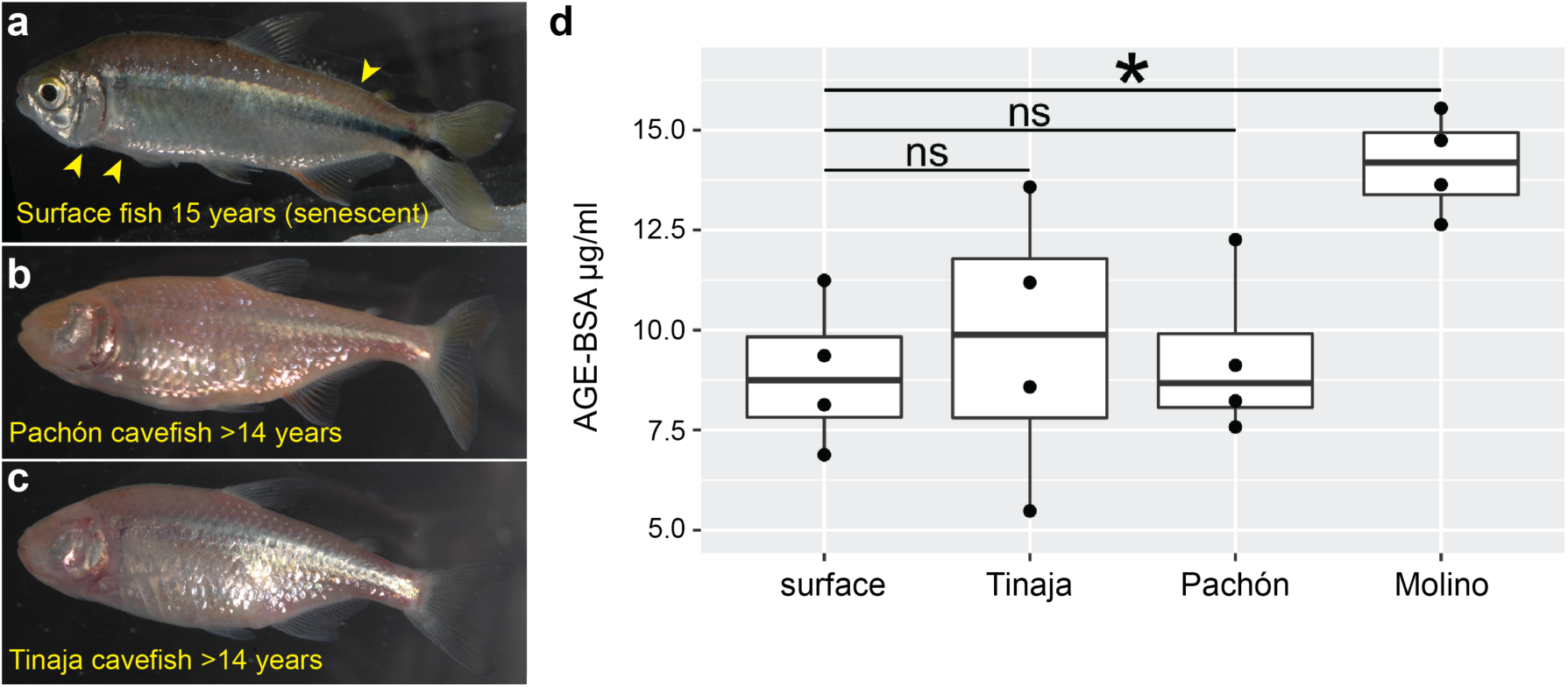
Despite elevated blood glucose levels and insulin resistance, Tinaja and Pachón cavefish do not show signs of senescence and do not accumulate advanced glycated end products in the blood. **a-c,** Surface fish (a), Pachón cavefish (b), and Tinaja cavefish (c) kept in the laboratory for the indicated duration and fed *ad libitum*. Cavefish were wild caught and the ages represent their minimum age. **a,** Surface fish shows signs of aging (yellow arrows) that are absent in cavefish at comparable ages (**b, c**). **d**, Quantification of advanced glycated end products (AGE-BSA g/mL) in serum from approximately 2-year-old surface, Tinaja, Pachón and Molino fish after a three-day fast (n = 4 for each population). Tinaja and Pachón AGE levels are not significantly different from surface (mean 9.7, 9.3, 8.9 respectively, ns, not significant). Molino AGE (mean 14) is significantly higher than surface (*p-value <0.05, one-way ANOVA with Tukey’s HSD post-hoc test).

Insulin-stimulated glucose uptake proceeds through phosphorylation of AKT at serine 473 (pAKT)^15^. We compared the ratio of pAKT vs AKT in freshly dissected cavefish and surface fish skeletal muscle treated with recombinant insulin (Fig. 1f, g). Consistent with the apparent dysregulation of glucose homeostasis, we observed lower pAKT levels in the Tinaja muscle after insulin treatment (mean 1.39 vs 0.775 average pAKT/AKT, p=0.017, Fig. 1f, g), suggesting that Tinaja cavefish are indeed insulin resistant relative to surface fish. To study if insulin resistance evolved as a common feature in derived cavefish populations, we tested insulin sensitivity in Pachón and Molino. Interestingly, Pachón muscle had lower pAKT levels in response to insulin (mean 0.806 average pAKT/AKT, p=.027, Fig. 1f, g), but despite having elevated blood glucose (Fig. 1b), Molino pAKT levels were equivalent to surface fish (mean 1.26 average pAKT/AKT, p=0.99, Fig. 1f, g). Our results suggest that Tinaja and Pachón evolved altered blood glucose regulation and insulin resistance in parallel, and that Molino may have evolved altered glucose metabolism through a different mechanism.

To gain insight into the genetic mechanism underlying the observed insulin resistance in cavefish, we examined the sequences of all known genes in the insulin pathway using the available genome sequence^16^ (Extended data 1). Strikingly, we found a coding difference in the insulin receptor gene (*insra*) between surface fish and cavefish, affecting a highly conserved proline in the extracellular cysteine rich domain (P211L, Fig. 2a-c). The presence of the mutation correlates with the insulin resistance phenotype, as both Tinaja and Pachón populations carried the mutation, while Molino cavefish showed the wildtype allele (Fig. 2b, c). Notably, the same genetic alteration is implicated in at least two known cases of Rabson-Mendenhall syndrome^17,18^, a form of severe insulin resistance in humans (Fig. 2c). The biochemical impact of the mutation has not been previously explored, but the position in the cysteine-rich domain suggests a role in insulin binding^19^. To test this, we generated transgenic HEK293T (Flp-In-293) cell lines that stably express the full-length surface fish or Tinaja cavefish *insra* and incubated the cells with different concenctrations of FITC-labeled human insulin. We measured fluorescence as a readout for binding efficiency using an image-based cytometry approach (ImagestreamX MarkII) and found that cells expressing the cavefish receptor displayed significantly lower binding at all but the lowest concentrations of insulin (Fig. 2d). Our results suggest that the *insra* P211L mutation affects insulin signaling by altering insulin binding efficiency.

We found that two independently-derived cavefish populations (from the Tinaja and Pachón caves) carry the same P211L mutation, suggesting parallel molecular evolution and selection for the mutation in the cave environment^21^. To extend these observations, we tested for the presence and frequency of the P211L mutation in fish living in their natural habitats. We sequenced 71 wild—caught surface fish samples from different localities, and 51 cavefish samples from 6 different caves (Extended data 2, Fig. 3a). In line with our previous observations, the mutation was absent in all the Molino samples (n=8), but present in all other tested cave populations (Tinaja, Yerbaniz, Pachón, Japonés, Arroyo, combined n=36). Notably, the cave populations carrying this mutation are all derived from the same ancestral stock of surface fish^22^. Although the mutation was partially present in heterozygote conditions in some of these caves, we did not find any cavefish that were homozygous for the surface allele despite continuous gene flow from the surface population^22^ (Fig. 3a). Our findings suggest the likelihood that there is active selection for the mutation in the caves, and also indicate a partially dominant effect of the cave allele. We did not detect the cave allele in any of the surface fish samples, suggesting that the mutation either appeared *de novo* in the cave populations, represents a rare variant not detected by our sampling frequency, or is absent in the current surface population, but was present in the ancestral surface fish stocks^22^. We reexamined the available whole genome data from different cave and surface populations (McGaugh et al, unpublished) to investigate these hypotheses. We found that *insr*a is not exceptional in its divergence between cave and surface fish using multiple independent statistical methods (F_ST_, D_XY_, of hapFLK results; Extended Data Table 1), arguing against a recent *de novo* occurrence of the mutation. In addition, we did not observe extreme reductions in diversity or exceptionally long tracts of homozygosity which would be indicative of recent hard selective sweeps (π, H-SCAN; Extended Data Table 1). Pachón cave, Tinaja cave, and surface populations exhibit negative Tajima’s D, which can under some scenarios indicate selection^23^, however, the values are not exceptional when compared to the rest of the genome. Our current hypothesis is that the mutation represents a rare variant in the ancestral or current stock of surface fish that fell under positive selection upon invasion of the caves.

We next sought to investigate the adaptive value of the mutation in the cave environment. Insulin resistance can result from obesity, but conversely it can also drive processes leading to adipogenesis and fat storage^24^. We found that cavefish weigh more than surface fish on a nutrient limited diet (2.08 vs 1.52 grams, p=0.02, Fig. 3b), raising the intriguing possibility that the insulin resistance in cavefish is part of an adaptive strategy to increase weight and thus survivability during food deprivation. To test for the influence of the *insra* mutation on weight, we genotyped and weighed 124 surface/Tinaja male F_2_ fish at approximately 1.5 years of age. We focused on males, as egg mass varies between individual females and can account for as much as 41% of female body weight (Extended Data Fig. 5), thus representing a potentially significant confounding variable if included in our analysis. Notably, we found that males carrying one or two copies of the cave P211L *insra* allele weigh on average 27% more than hybrids carrying only the surface allele (1.63 vs 1.28 grams, p = 0.006, Fig. 3c). To test if the difference in weight is entirely attributable to metabolic differences in processing ingested food or is also affected by differences in consumption when fed *ad libitum*, we individually housed the fish and fed them a controlled diet of 6 mg of food per day for 4 months. On average, both carriers and non-carriers of the mutated *insra* allele gained weight, but no significant difference was observed in percent weight gain (17.3% and 20.5%, respectively; p-value: 0.68), underscoring the importance of overeating in combination with insulin resistance to increase body weight^8^. After four months on the controlled diet, carriers of the cave allele still weighed significantly more than homozygous surface carriers (1.82 vs 1.47 grams, p=0.004, Fig. 3d). These findings indicate a role of the cavefish *insra* locus on weight gain, but cannot entirely exclude the possibility of another effect of a different gene in cis to the P211L mutation.

To verify that the phenotypes we have associated with the P211L mutation in cave fish are indeed due to the alteration of the *insra* gene, we used CRISPR gene editing to introduce the P211L mutation into the *insra* gene in zebrafish (*Danio rerio*) via homology directed repair^25^ (Supp. Figure 1, Figure 3e). We have shown that the P211L mutation causes a decrease in insulin binding (Fig. 2d). To determine if decreased binding causes insulin resistance in zebrafish, we measured the ratio of pAKT/AKT in freshly dissected zebrafish skeletal muscle treated *ex vivo* with insulin or PBS (Fig. 3f). We found that the ratio of pAKT/AKT is less in zebrafish that are homozygous for the P211L mutation compared to heterozygous fish for both PBS (0.05 vs 0.13, *p=0.016) and insulin treated conditions (0.13 vs 0.32, p=0.067, n=3 fish per genotype and condition). We next tested whether this insulin resistence is sufficient to explain the increase in weight we observe in cavefish carrying the P211L mutation. We paired heterozygous fish with successful germline transmission and weighed their progeny shortly before maturity (55 dpf) to avoid gonadal effects on weight. We found that zebrafish homozygous for the cave allele are longer (20.0 mm vs 18.4 mm, p=0.0046) and weigh more (124.6 mg vs 99.7 mg, p=0.022) than their siblings raised under the same conditions (Fig. 3e, g, h). Thus, our findings show that the P211L mutation is sufficient to explain both the insulin resistance and increased weight that we observe in Tinaja cavefish.

The extreme metabolic phenotypes, reminiscent of Type 2 diabetes (insulin resistance, hyperglycemia), in conjunction with the fact that cavefish have a fatty liver^8^, raise the question whether cavefish have pathologies that are characteristic of these phenotypes in other species and are therefore less healthy than surface fish. If so, it would suggest an evolutionary trade-off wherein the cavefish might have sacrificed other aspects of physiological health to reap the benefits of starvation resistance. Alternatively, the fish could have evolved unique compensatory mechanisms allowing them to remain healthy despite potential deleterious metabolic changes. In support of the latter idea, surface fish and cavefish live longer than fifteen years in the laboratory with no noticeable differences in fertility decline. However, we find that surface fish begin to exhibit classic signs of aging in fish (e.g. ^27^) by age fifteen, such as sunken skin, tattered fins, and a hunched back (Fig 4a) while Tinaja and Pachón can live in excess of fourteen years without these indications of senescence (Fig. 4b.c).

In humans, sustained high blood glucose levels and dysregulation of glucose homeostasis can lead to cell damage over time. A major cause of morbidity in human diabetic patients with long-term elevation in blood glucose is tissue damage caused by excessive non-enzymatic glycation of proteins in the blood, generating advanced glycation end-products (AGEs)^28^. AGEs are closely associated with diabetes induced vascular damage, cardiovascular disease, and aging^29^. We compared the level of AGEs in the serum of two year-old cavefish and surface fish that had been fed *ad libitum* their entire lives (Fig. 4b). Interestingly, we did not detect any differences in the levels of AGEs between Tinaja and Pachón cavefish, relative to surface fish (mean of 9.7, 9.3, and 8.9 µg/ml respectively, p=0.99, 0.95), in spite of elevated blood glucose levels in fish from these cave populations (Fig. 1b). This result suggests that these two cavefish populations may have mechanisms for reducing protein glycation, rendering them impervious to the damaging effects of elevated blood glucose. Notably, the Molino cavefish, which we found are not insulin resistant but nonetheless show higher blood sugar levels, do have elevated levels of AGEs compared to surface fish (mean 14.1 vs. 8.9 µg/ml, p=0.03). It remains to be determined if the health and longevity of the Molino population is influenced by accumulation of AGEs, but our results suggest they may have evolved altered blood glucose homestasis through a different mechanism than Tinaja and Pachón cavefish. Insulin resistance in human patients carrying the Tinaja and Pachón-specific mutation in *insra* is accompanied by additional sequelae, including dental dysplasia, and growth retardation^17,18^, neither of which are observed in cavefish^30^.

All organisms need to adapt to the environments in which they live. This includes optimizing the utilization of the nutrients available to them, in addition to evolving strategies for finding food and for avoiding predation. Yet far less is known about metabolic adaptation than, for example, morphological evolution. Our results suggest a unique, and perhaps surprising, metabolic strategy adopted by cavefish in response to annual extended periods of extreme nutrient deprivation. In particular, the cave morphs of *Astyanax mexicanus* alter blood glucose levels as a partial mechanism for rapidly increasing body weight during brief periods of abundance. These findings highlight the extreme physiological measures that can evolve in critical, and otherwise highly conserved, metabolic pathways to accommodate exceptional environmental challenges. Additionally, our findings establish cavefish as a natural model to investigate resistance to pathologies of diabetes-like dysregulation of glucose homestasis. As such, further understanding cavefish metabolism may reveal factors that can help eliminate both short-term and long-term negative effects of elevated blood glucose levels in humans, as well as providing further insight into the metabolic changes that can allow organisms to adapt to extreme environments.

## Methods

### Fish husbandry and diet

Unless stated otherwise, fish were fed *ad libitum* with a combination of New Life Spectrum TheraA+ small fish formula and *Artemia,* and housed at densities of less than, or equal to, two adult fish per liter of water. F2 hybrids were housed individually in 1.5L tanks and fed three pellets (∼6mg) of New Life Spectrum TheraA+ small fish formula once per day for >4 months. For the starvation experiment, fish were moved to individual containers and water was changed daily.

### Blood glucose, glucose tolerance, and arginine tolerance

Blood was collected from the caudal tail vein using a U-100 insulin needle and glucose was measured using Freestyle lite blood glucose meter and test strips. Glucose (2.5mg/gram fish), arginine (6.6µM/gram fish), or PBS was injected into the intraperitoneal cavity using a U-100 insulin needle.

### pAKT quantification

We quantified pAKT level in fillets of skeletal muscle taken directly after fish decapitation. For *A. mexicanus,* skeletal muscle was cut into three equal strips per fish. Strips were incubated in PBS, X, 0.1X concentration of recombinant Human insulin for 25 mins (sigma product I9278, X = 9.5-11.5 μg/mL insulin). The tissues were rinsed in pbs and then homogenized and lysed in RIPA buffer (SIGMA) with protease and phosphatase inhibitor (Pierce^(tm)^ Protease and Phosphatase Inhibitor Mini Tablets, EDTA Free) for 30 mins. Protein concentration was measured via BCA (Pierce). Lysate protein concentrations were then equalized and run on 4-12% Bis-Tris protein gel and transferred on a nitrocellose membrane. Blots were probed for AKT (Cell Signalling). Following stripping, blots were probed for phospho-AKT (ser473) (Cell Signalling). Densitometry was done on imageJ. For *D. rerio* two fillets of skeletal muscle were removed from both sides of fish directly after decapitation. Fillets were rinsed in PBS and then incubated in PBS or 10µg/mL human recombinant insulin (sigm, I0908) for approximatley 40 minutes. The skin was then removed from the skeletal muscle and the muscle was finley minced using a scalpel. We quantified the ratio of pAKT/AKT using the Akt(pS473) + total Akt ELISA Kit (abcam ab126433) according to the manufacturers protocol. We used 200µL lysis buffer per sample and loaded 85µL of lysate per well.

### INSRA P211L genotyping

Genomic DNA from tail fin clips was diluted 5-fold and used as target DNA to amplify the *insra* locus using the following oligonucleotide primers: insra_f: GCACCCTTACACCCTTACATGA; insra_r: TACCGCTCAGCACTAATTTGGA; Product size: 700 bp. PCR reactions were carried out in 12.5 μl volume containing 1X LA PCR Buffer II (Clontech), 2.5 mM MgCl2, 0.4 mM dNTP mix, 0.4 uM of each forward and reverse primer and 0.05 units of TaKaRa LA Taq DNA Polymerase (Clontech). The PCR cycling conditions were as follows: Initial denaturation at 94°C for 2 min, followed by 35 cycles of 94°C for 30 sec, annealing temperature 52°C for 30 sec and 72°C for 1 min. A final 5 min elongation step was performed at 72°C. The PCR products were diluted 10-fold and sequenced directly on a 3730XL DNA Analyzer (Applied Biosystems) using the sequencing primer: GGTGGAGTTGATGGTGGTATAG.

### Selection scans at the *insra* locus

We examined the insra locus with data that is a part of an ongoing genome-wide selection and demography companion study (Herman et al. in preparation). Methods are explained in greater detail in the companion study, but briefly, we used Illumina Hiseq 2000 to sequence 100bp reads from 6-10 individuals from each population of Tinaja cave, Molino cave, Pachón cave, Rascon surface, and Río Choy surface populations, (total N = 43) and two individuals from the sister taxa *A. aeneus*. Individuals were sequenced with v3 chemistry. Reads were cleaned with Trimmomatic v0.30^31^ and cut-adapt v1.2.1 (http://dx.doi.org/10.14806/ej.17.1.200) and aligned to the reference Pachón genome using bwa-mem algorithm in bwa-0.7.1^32^ resulting in an aligned coverage depth of ∼7-12x. Variants were called using the Genome Analysis Toolkit v.3.3.0 (GATK)^33^ and Picard v1.83 (http://broadinstitute.github.io/picard/). Outlier scan metrics (π, DXY, FST, Tajima’s D) were conducted using VCFtools v0.1.13^34^ and custom scripts. HSCAN (http://messerlab.org/software/) and hapFLK^35^ were also used in examining *insra* for outliers. Metrics were dense ranked across the genome and the ranking position of *insra* was used to determine if it was exceptionally divergent between cave and surface populations relative to the rest of the genome.

### Glucagon and Insulin quantification

The number of cells producing insulin or glucagon were determined using the following protocol: 10-11dpf fish were euthanized with an overdose of tricane and fixed in 4% paraformaldehyde overnight at 4°C. Fish were washed in PBST, transferred to water for 1 minute, acetone for 10 minutes at −20?, water for 1 minute, then washed in PBST. Blocking in 5% donkey serum, 1% DMSO, and 0.2% BSA was done for 1 hour at room temperature. Fish were incubated with primary antibodies (1:200 sheep anti-Glucagon (abcam), 1:200 guinea pig anti-Insulin (Dako)) and then secondary antibodies (1:400 donkey anti-sheep 488, 1:400 goat anti-guinea pig 647) overnight at room temperature in glass vials, then washed with PBST, stained with DAPI, and imaged. Images were collected at 63X using a 1.0µm z-stack on a Zeiss 780 confocal microscope. Nuclei surrounded by insulin or only glucagon were counted manually using FIJI cell counter.

To quantify circulating glucagon level, we collected serum from the caudal tail vein of fish that were approximately 2-years-old and were fasted for 24 hours (n=12 for each population). The serum was used for a Glucagon radioimmunoassay according to the manufacturer protocol (MGL-32K; Millipore, Billerica, MA). To quantify circulating insulin level we collected plasma from the caudal tail vein of 2-year-old and 1-year-old fish and blotted the serum onto a nitrocellulose membrane using a BIO-RAD Bio-dot sf device. Ponceau protein staining was used to verify equal protein loading after which insulin was probed using anti—Insulin (DAKO). Quantification of insulin levels was done using FIJI-imageJ.

### Insulin Binding Experiment

The full-length *Astyanax mexicanus insra* protein-coding sequence was amplified from Surface fish (S) and Tinaja cavefish (T) cDNA and cloned into a modified pcDNA3.1/Hygro vector providing an N-terminal FLAG epitope tag^36^. To generate stable cell lines, the FLAG-tagged *insra* cassettes were cloned into pcDNA5/FRT vector (Invitrogen cat# V601020) allowing for Flp recombinase—mediated integration into the Flp-In-293 cell line (Invitrogen cat# R75007) per the manufacturer’s procedures. One positive clone from each S and T cell line was selected and used for the insulin-binding assays. 100mm plates were seeded at 30% confluency and cultured in DMEM/10%FBS+1xGlx media for 48 hours. The plates were then changed to insulin-free FreeStyle 293 Expression Medium (cat# 12338018) and incubated for an additional 24 hours. Plates at ∼70-80% confluency were pre-chilled for 30 min at 4°C, and the medium replaced with 5 ml of cold FreeStyle 293 Expression Medium containing 42mM HEPES pH7.5 and human FITC labeled insulin (Sigma cat# I3661) at final concentrations of 0, 0.1, 0.2, 0.4, 0.8, 1 and 3 µM. For binding competition, 10 µM of unlabeled human insulin (Sigma cat# I9278) and 1 µM of human FITC labeled insulin were added. After one hour incubation in the dark at 4°C, the medium was removed and the plates were washed with 5mL of cold 1x PBS. Cells were dissociated in 2mL of 0.5mM EDTA in PBS at 37°C for 7 min, transferred into Eppendorf tubes, pelleted for 5 min at 200xg at 4°C and resuspended in 1mL of cold 1x PBS. To stain dead cells, 1 μl of Fixable Viability Dye (FVD eFluor450, Invitrogen cat# 65-0863) was added to each 1mL of cell suspension and incubated on ice in the dark for 20 min. Subsequently, the cells were washed once with cold 1x PBS and fixed in 1mL of 4% formaldehyde. After two more washes with cold 1x PBS, the cells were resuspended in 150μl of cold 1x PBS, filtered through a 70μm cell strainer (Filcons 070-67-S) and transferred into a round-bottom 96-well plate. Binding data was acquired on an ImagestreamX MarkII (EMD Millipore) at 40x. FITC was excited with 150mW 488nm on camera 1 and detected on channel 2. Fixable live/dead was excited with 12mW 405nm on camera 2 and detected on channel 7. Single color controls were used for color compensation. Bright field was collected on channels 5 and 11. Analysis was performed in IDEAS v6.2 and fluorescence intensity was reported as integrated intensity within an adaptive erode mask for bright field.

### Genome editing in zebrafish using the CRISPR/Cas9 technology

#### Guide-RNA and donor design

We designed the guideRNA target sites using the web tool by the MIT Zhang lab (http://crispr.mit.edu). We then validated the target region and checked for SNPs by PCR and sequencing of genomic DNA. We used Cas9 protein from PNABio and 2-part Alt-R guideRNAs from IDT. Single-stranded oligodeoxynucleotides (ssODNs) were ordered as Ultramers from IDT for generating the SNP mutations. The ssODNs included 100bp of homology arms as well as silent mutations in the guideRNA target site to prevent re-annealing of the guideRNA following homologous recombination. We also protected both ends of the ssODNs with 3 phosphorothioate bonds to inhibit exonuclease degradation in the cell.

#### Microinjection

We annealed the specific crRNA with tracrRNA to form the guideRNA complex, followed by hybridizing with the Cas9 protein to form a ribonucleoprotein (RNP) complex. The RNP was then injected into at least 200 and up to 1500 zebrafish embryos at the 1-cell stage.

#### Screening and breeding

We designed genotypic screening assays to find mutants and confirm the expected location of the mutation. After injections, we screened for mutation success rate in the F0. We then outcrossed mosaic individuals to wildtype and tested for germline transmission rates and to raise potential heterozygous mutants.

### Quantification of Advanced Glycation Endproduct (AGEs)

We used Oxiselect^(tm)^ Advanced Glycation End Product (AGE) Competitive Elisa Kit (San Diego, CA) according to the manufacturers protocol to measure AGE level in serum from 2-year-old fish that were fasted for 3 days.

### Statistics and Figure preparation

Figures and statistics were produced using R^37^ and ggplot2^38^ package.

### Data availability

The datasets generated during the current study are available in the Stowers original data repository and/or available from the corresponding author on reasonable request.

## Acknowledgments

We thank Yashodhan Chinchore and Cem Sengel for technical advice; Xin Gao for bioinformatic support; Zachary Zakibe for photographs of the fish; the entire Aquatics facility at Stowers for fish maintenance and support; the cell culture core at Stowers for cell line maintenance and advice; the molecular biology core at Stowers for design, execution, and validation of the CRISPR constructs; the proteomics core; in particular, Michaella Levy for advice and computational modeling of the insulin receptor; Adam Herman for help with the genome scan; Mark Miller for illustration; and Stacey Williams, Fleur Damen, and Kira Fox for helpful feedback on the manuscript text. This work was supported by a grant from the NIH to C.J.T. (HD089934) and institutional funding to N.R. M.R.R was supported by a National Research Service Award (DK108495).

## Author contributions

Author contributions: AA, MR, NR, CT conceived of project and designed research with additional contributions from KG, RP, AB. AA, MR, KG, RP, JS, BM, MP, AB, JT, SM, RB, NR performed the research. AA, MR, CT, NR wrote the paper.

## Author information

The authors declare no competing financial interest. Correspondence and request for materials should be addressed to

## References

1 Staples, J. F. Metabolic Flexibility: Hibernation, Torpor, and Estivation. Compr Physiol 6, 737–771 (2016).

2 Culver, D. C. & Pipan, T. The Biology of Caves and Other Subterranean Habitats. (Oxford University Press, 2009).

3 Gross, J. B. The complex origin of Astyanax cavefish. BMC Evol Biol 12, 105 (2012).

4 Bradic, M., Teotonio, H. & Borowsky, R. L. The population genomics of repeated evolution in the blind cavefish Astyanax mexicanus. Mol Biol Evol 30, 2383–2400 (2013).

5 Carlson, B. M., Onusko, S. W. & Gross, J. B. A high-density linkage map for Astyanax mexicanus using genotyping-by-sequencing technology. G3 (Bethesda) 5, 241–251 (2015).

6 Elipot, Y., Legendre, L., Pere, S., Sohm, F. & Retaux, S. Astyanax transgenesis and husbandry: how cavefish enters the laboratory. Zebrafish 11, 291–299 (2014).

7 Casane, D. & Retaux, S. Evolutionary Genetics of the Cavefish Astyanax mexicanus. Adv Genet 95, 117–159 (2016).

8 Aspiras, A. C., Rohner, N., Martineau, B., Borowsky, R. L. & Tabin, C. J. Melanocortin 4 receptor mutations contribute to the adaptation of cavefish to nutrient-poor conditions. Proc Natl Acad Sci U S A 112, 9668–9673 (2015).

9 Moran, D., Softley, R. & Warrant, E. J. Eyeless Mexican cavefish save energy by eliminating the circadian rhythm in metabolism. PLoS One 9, e107877 (2014).

10 Hüppop, K. Oxygen consumption of Astyanax fasciatus (Characidae, Pisces). A comparison of epigean and hypogean populations. Environ. Biol. Fishes 17, 299–308 (1986).

11 Penney, C. C. & Volkoff, H. Peripheral injections of cholecystokinin, apelin, ghrelin and orexin in cavefish (Astyanax fasciatus mexicanus): effects on feeding and on the brain expression levels of tyrosine hydroxylase, mechanistic target of rapamycin and appetite-related hormones. Gen Comp Endocrinol 196, 34–40 (2014).

12 Saltiel, A. R. & Kahn, C. R. Insulin signalling and the regulation of glucose and lipid metabolism. Nature 414, 799–806 (2001).

13 Rines, A. K., Sharabi, K., Tavares, C. D. & Puigserver, P. Targeting hepatic glucose metabolism in the treatment of type 2 diabetes. Nat. Rev. Drug Discov. 15, 786–804 (2016).

14 Navarro I, e. a. Insights into Insulin and Glucagon Responses in Fish. Fish Physiology and Biochemistry 27, 205–216 (2002).

15 Lizcano, J. M. & Alessi, D. R. The insulin signalling pathway. Curr Biol 12, R236–238 (2002).

16 McGaugh, S. E. et al. The cavefish genome reveals candidate genes for eye loss. Nat Commun 5, 5307 (2014).

17 A. Atray, S. J., K. Thai, P. Hiremath, R.M. Anjana, R. Unnikrishnan, V. Mohan, V. Radha. Rabson Mendenhall Syndrome; a case report. Journal of Diabetology 2 (2013).

18 Carrera, P. et al. Substitution of Leu for Pro-193 in the insulin receptor in a patient with a genetic form of severe insulin resistance. Hum Mol Genet 2, 1437–1441 (1993).

19 Taylor, S. I. et al. Mutations in insulin-receptor gene in insulin-resistant patients. Diabetes Care 13, 257–279 (1990).

20 De Meyts, P. & Whittaker, J. Structural biology of insulin and IGF1 receptors: implications for drug design. Nat. Rev. Drug Discov. 1, 769–783 (2002).

21 Wood, T. E., Burke, J. M. & Rieseberg, L. H. Parallel genotypic adaptation: when evolution repeats itself. Genetica 123, 157–170 (2005).

22 Bradic, M., Beerli, P., Garcia-de Leon, F. J., Esquivel-Bobadilla, S. & Borowsky, R. L. Gene flow and population structure in the Mexican blind cavefish complex (Astyanax mexicanus). BMC Evol Biol 12, 9 (2012).

23 Tajima, F. Statistical method for testing the neutral mutation hypothesis by DNA polymorphism. Genetics 123, 585–595 (1989).

24 Kahn, B. B. & Flier, J. S. Obesity and insulin resistance. J. Clin. Invest. 106, 473–481 (2000).

25 Albadri, S., Del Bene, F. & Revenu, C. Genome editing using CRISPR/Cas9-based knock-in approaches in zebrafish. Methods 121-122, 77–85 (2017).

26 Borowsky, R. Restoring sight in blind cavefish. Curr Biol 18, R23–24 (2008).

27 Hayes, A. J. et al. Spinal deformity in aged zebrafish is accompanied by degenerative changes to their vertebrae that resemble osteoarthritis. PLoS One 8, e75787 (2013).

28 Yan, S. F., Ramasamy, R. & Schmidt, A. M. Mechanisms of disease: advanced glycation end-products and their receptor in inflammation and diabetes complications. Nat. Clin. Pract. Endocrinol. Metab. 4, 285–293 (2008).

29 Prasad, A., Bekker, P. & Tsimikas, S. Advanced glycation end products and diabetic cardiovascular disease. Cardiol. Rev. 20, 177–183 (2012).

30 Keene, A., Yoshizawa, M. & McGaugh, S. Biology and Evolution of the Mexican Cavefish. (New York: Academic Press, 2015).

31 Bolger, A. M., Lohse, M. & Usadel, B. Trimmomatic: a flexible trimmer for Illumina sequence data. Bioinformatics 30, 2114–2120 (2014).

32 Li, H. & Durbin, R. Fast and accurate long-read alignment with Burrows-Wheeler transform. Bioinformatics 26, 589–595 (2010).

33 Van der Auwera, G. A. et al. From FastQ data to high confidence variant calls: the Genome Analysis Toolkit best practices pipeline. Curr Protoc Bioinformatics 43, 11 10 11–33 (2013).

34 Danecek, P. et al. The variant call format and VCFtools. Bioinformatics 27, 2156–2158 (2011).

35 Fariello, M. I., Boitard, S., Naya, H., SanCristobal, M. & Servin, B. Detecting signatures of selection through haplotype differentiation among hierarchically structured populations. Genetics 193, 929–941 (2013).

36 Tomomori-Sato, C. et al. A mammalian mediator subunit that shares properties with Saccharomyces cerevisiae mediator subunit Cse2. J Biol Chem 279, 5846–5851 (2004).

37 Team, R. C. R: A language and environment for statistical computing, <https://www.R-project.org/>. (2016).

38 Wickham, H. ggplot2: Elegant Graphics for Data Analysis. (Springer-Verlag, New York, 2009).

